## Supplementary Material for "GENIUS: GEnome traNsformatIon and spatial representation of mUltiomicS data"

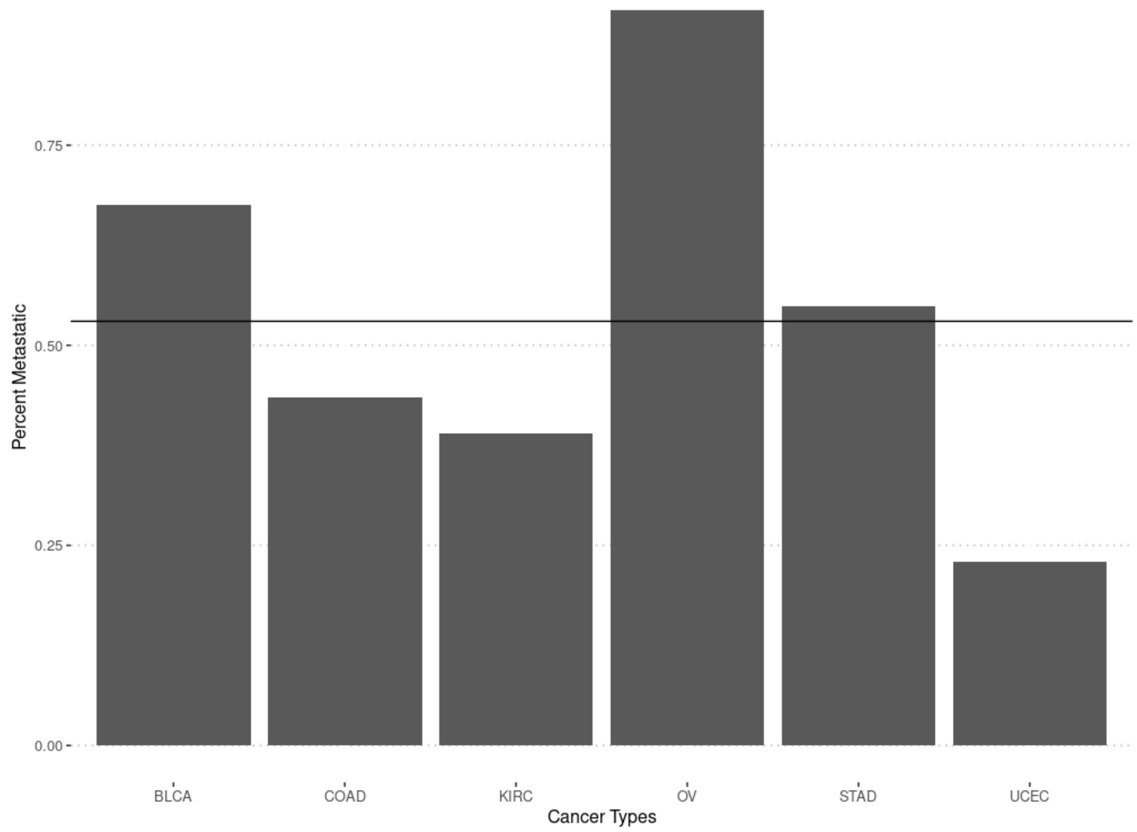


**Supplementary Figure 1:** Figure showing percent of sample classified as metastatic when using stage as variable.

A

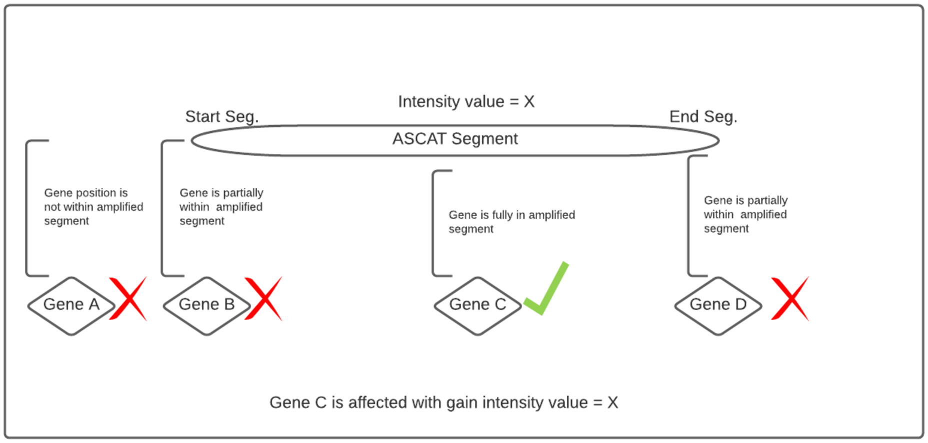


B

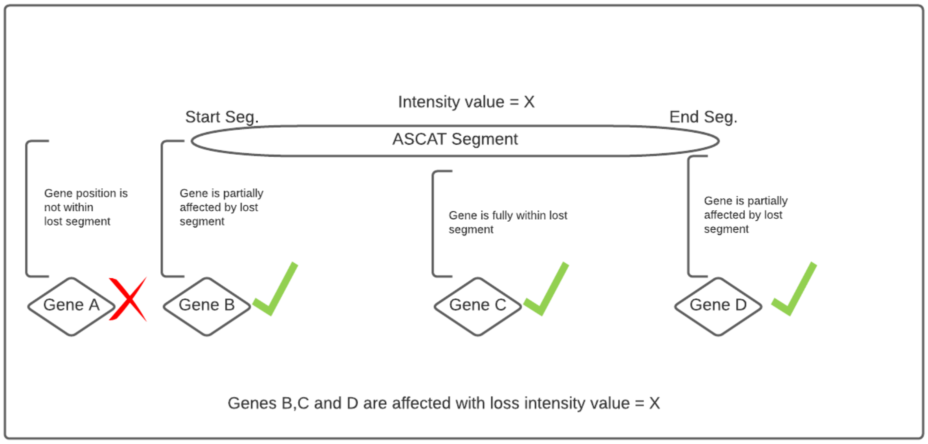


**Supplementary Figure 2:** Overall representation of a process where chromosomal instability information was included in genome image construction. (A) Amplification of a gene was declared when the gene was fully within an amplified segment. (B) Deletion of a gene was declared when the gene was partially within a deleted segment.

A

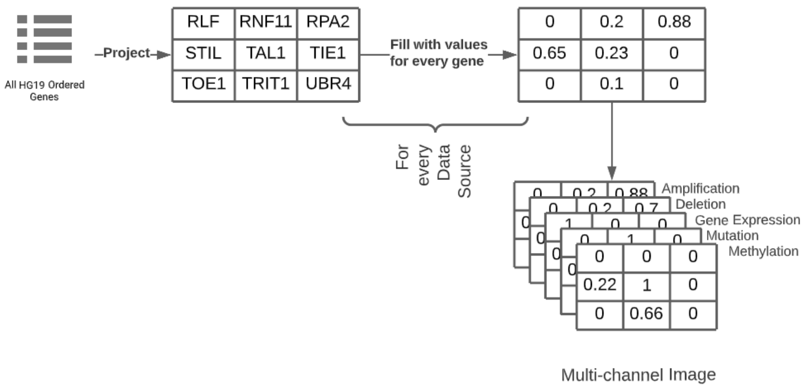


B

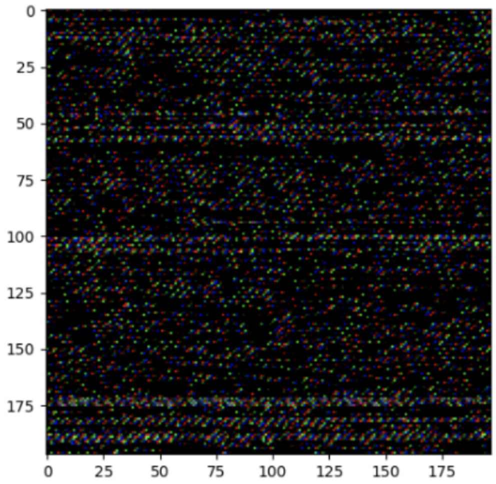


**Supplementary Figure 3:** Figure representing an example of genome image construction. (A) HG19 genes were ordered by chromosome and chromosome position. Next, based on gene position, the matrix was filled, where each cell represents a single gene. This was used for a template when creating matrices for every data source included in genome image construction. (B) Example of genome image.

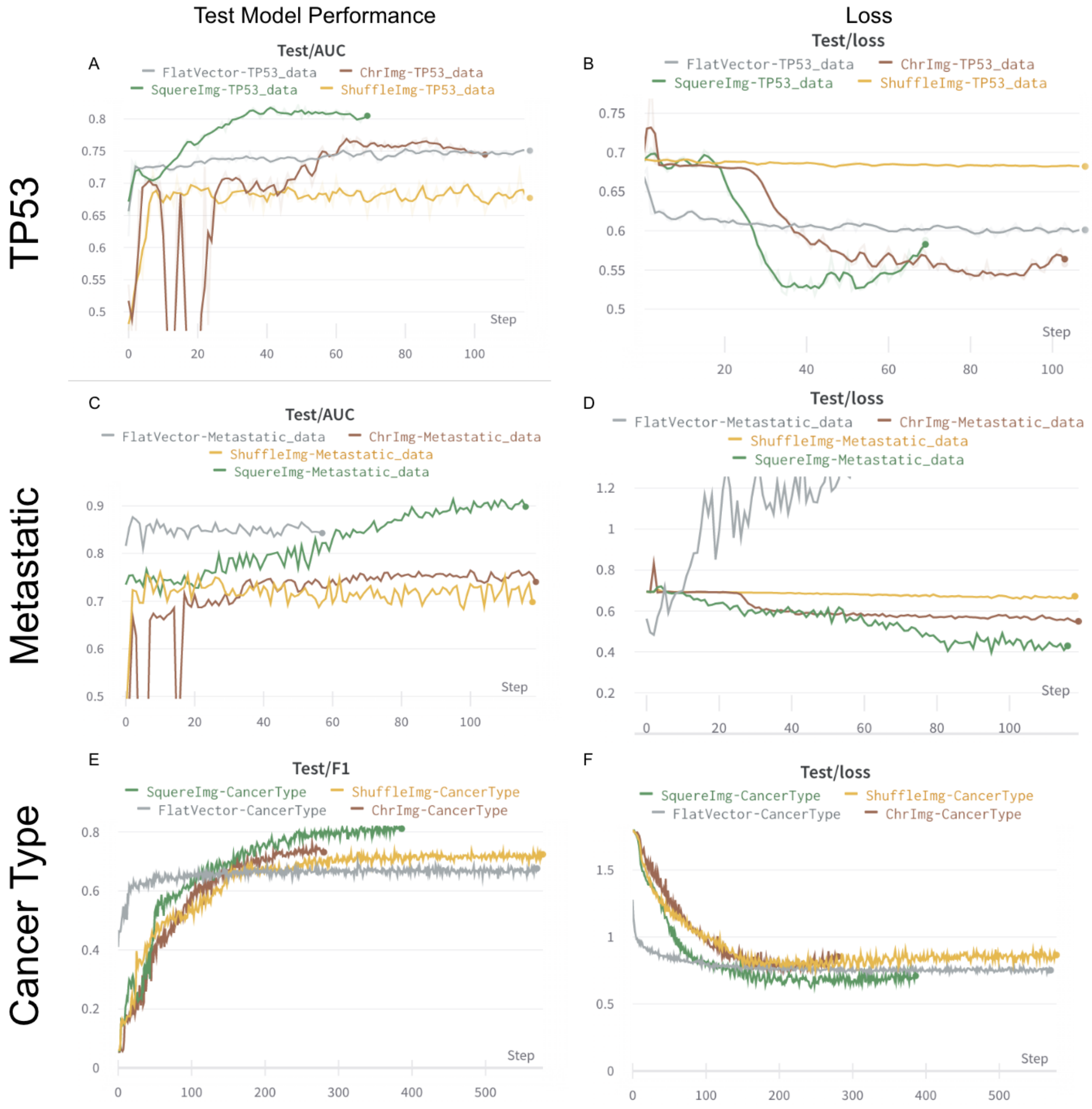


**Supplementary Figure 4:** Figure showing the process of training four image transformations in classification scenarios. X-axis represents epochs and Y-axis represents AUC/F1 (Panels A,C,E). On panels B, D, F, Y-axis is switched with loss.

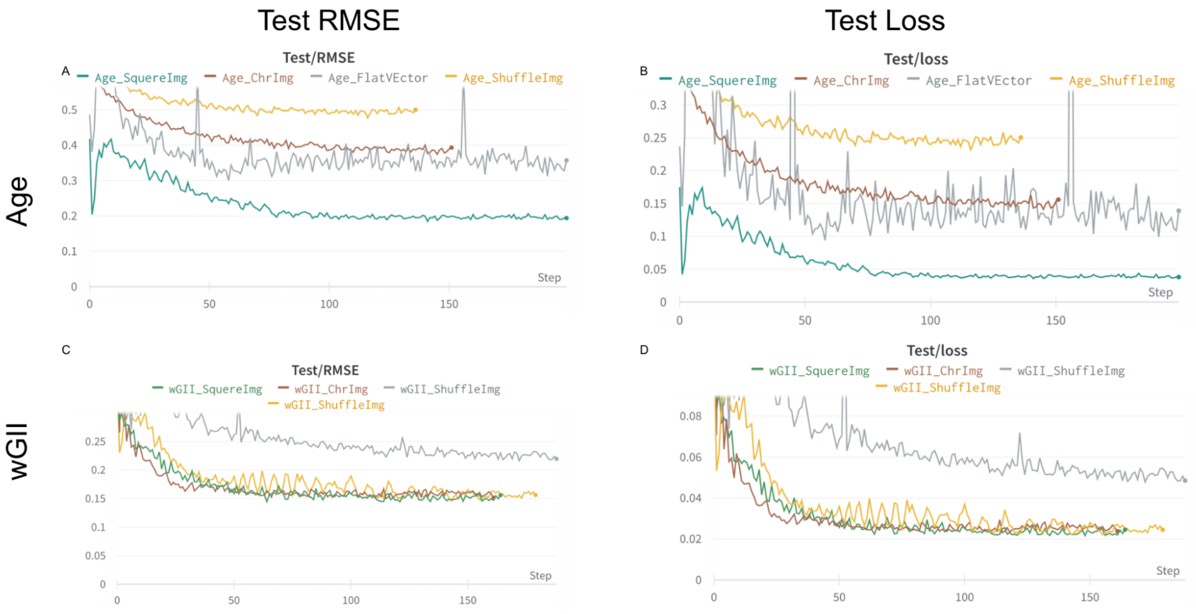


**Supplementary Figure 5:** Figure showing the process of training four image transformations in regression scenarios. X-axis represents epochs and Y-axis represents RMSE.

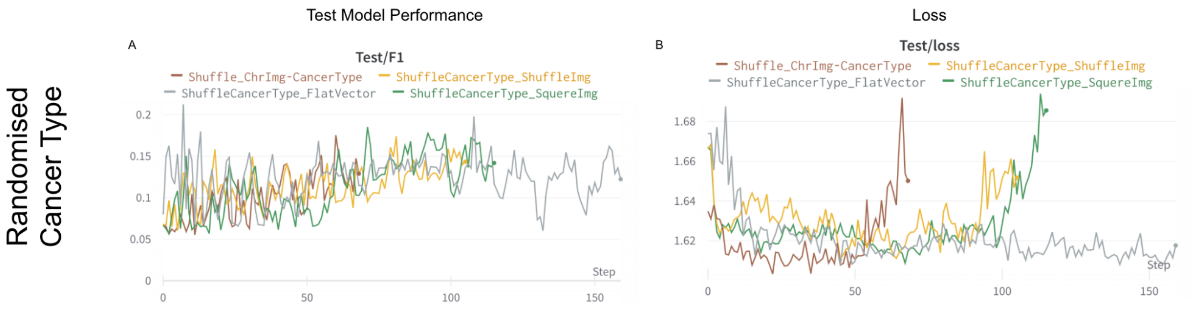


**Supplementary Figure 6:** Figure showing the process of training four image transformations in classification scenarios (negative control). X-axis represents epochs and Y-axis represents F1 (Panels A). On panel B, Y-axis is switched with loss.

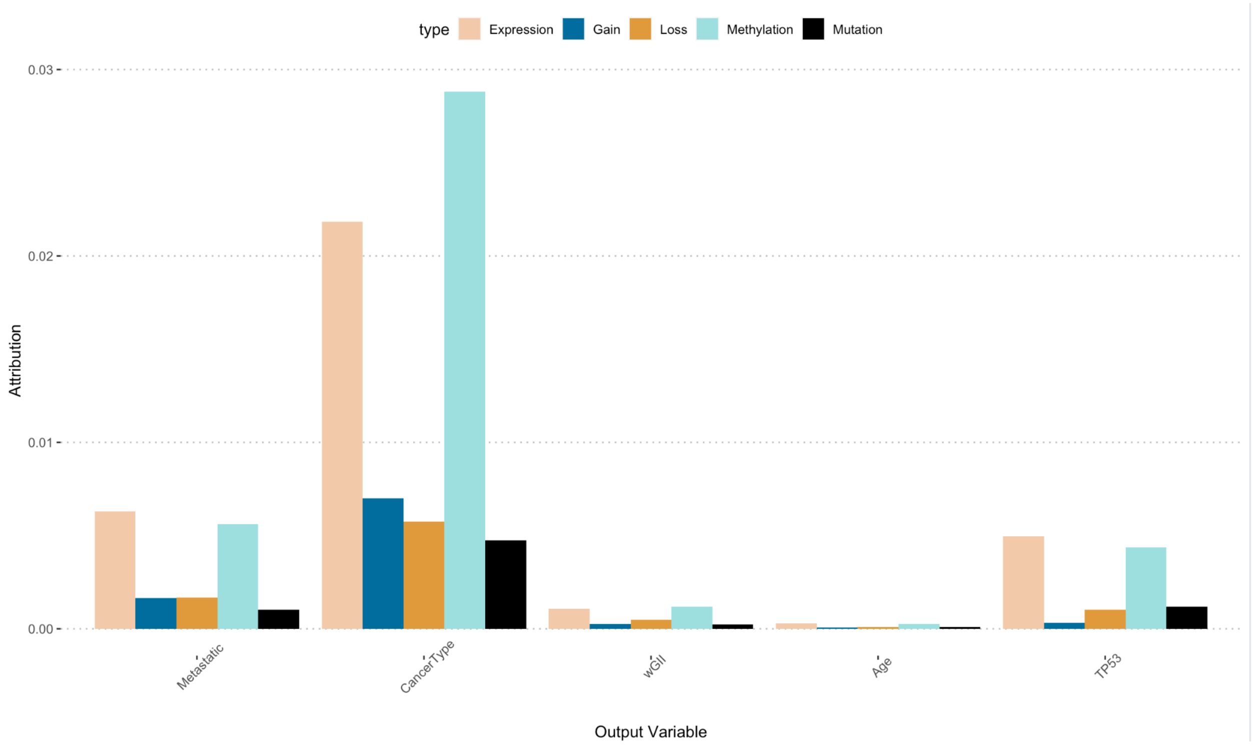


**Supplementary Figure 7:** The figure shows the relative attribution of each variable we predicted using the GENIUS framework. The color represents the relative importance of a specific data source in a specific scenario (x-axis).

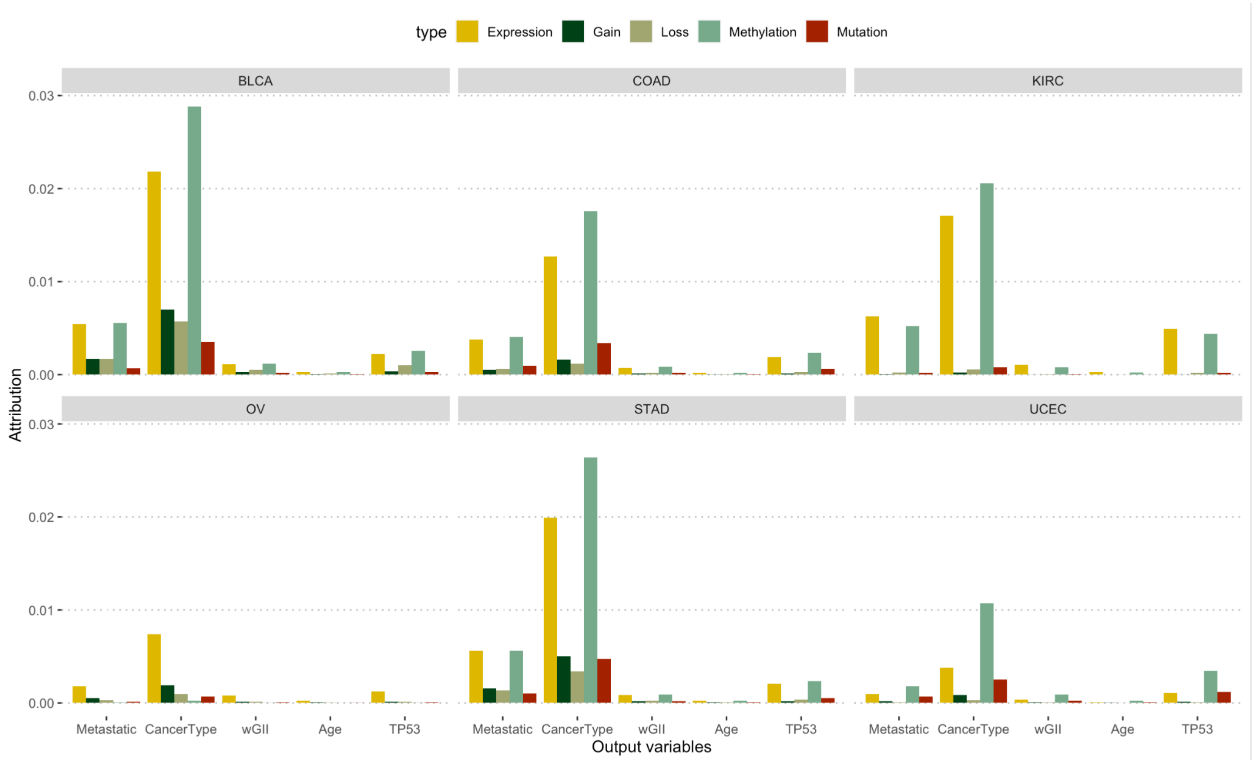


**Supplementary Figure 8:** The figure shows the relative attribution of each variable we predicted using the GENIUS framework (x-axis) but split by cancer type. The color represents the relative importance of a specific data source in a specific scenario (x-axis) and each panel represents cancer type.

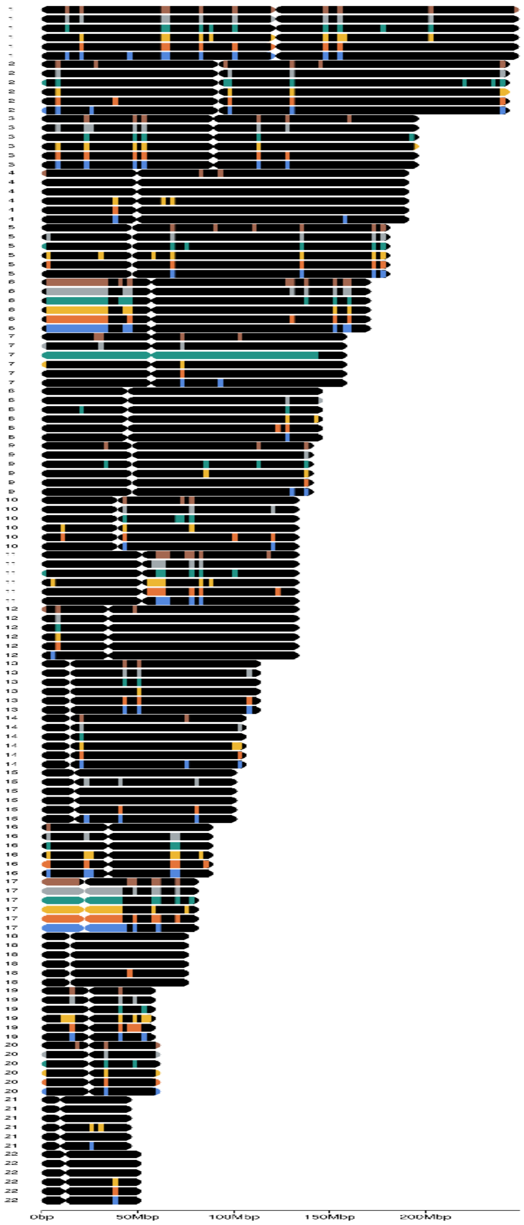


**Supplementary Figure 9:** The figure shows patterns of methylation across chromosomes 1-22.

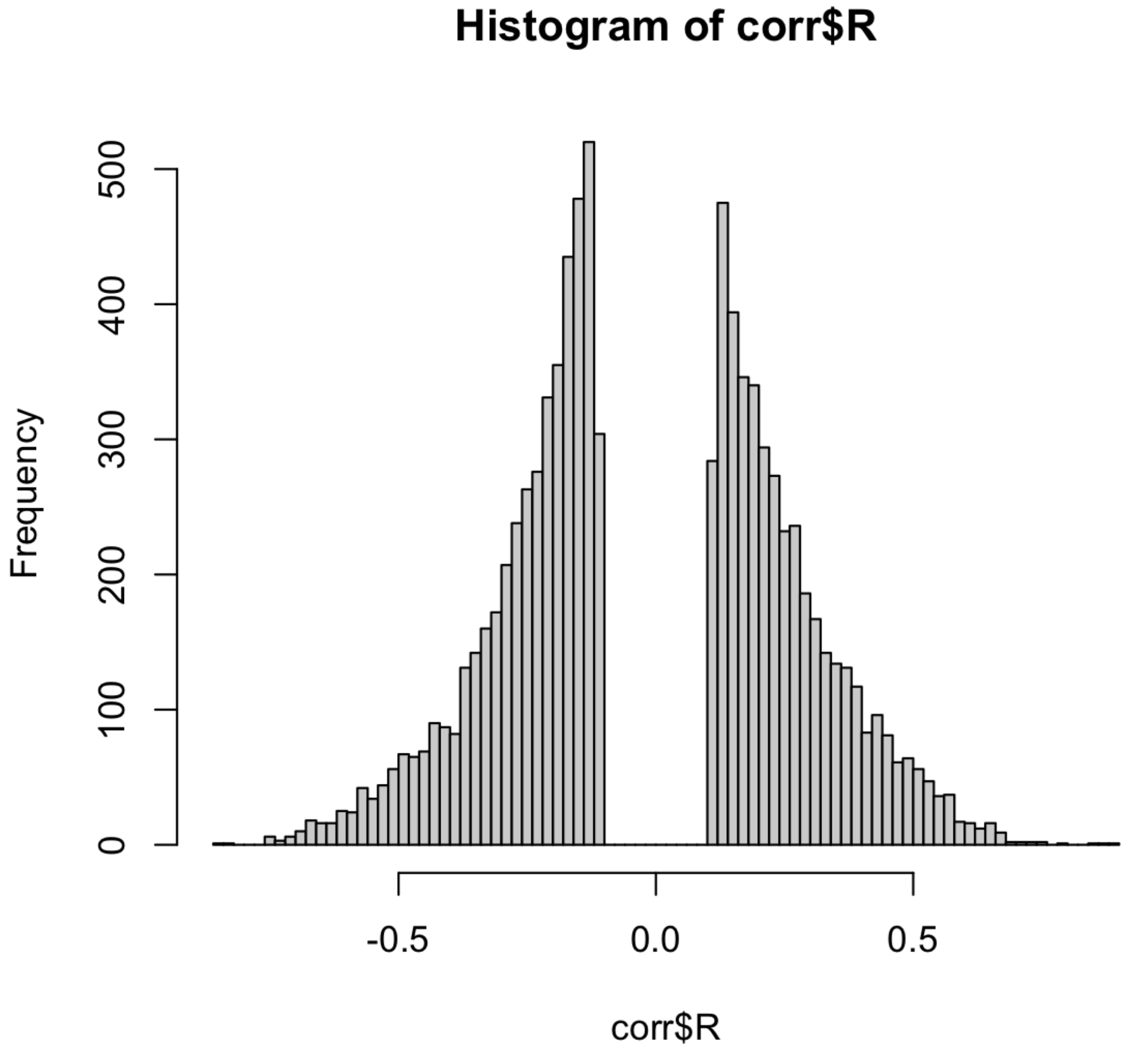


**Supplementary Figure 10:** Histogram of correlation coefficients between each gene found in BLCA TCGA gene expression data and methylation data.

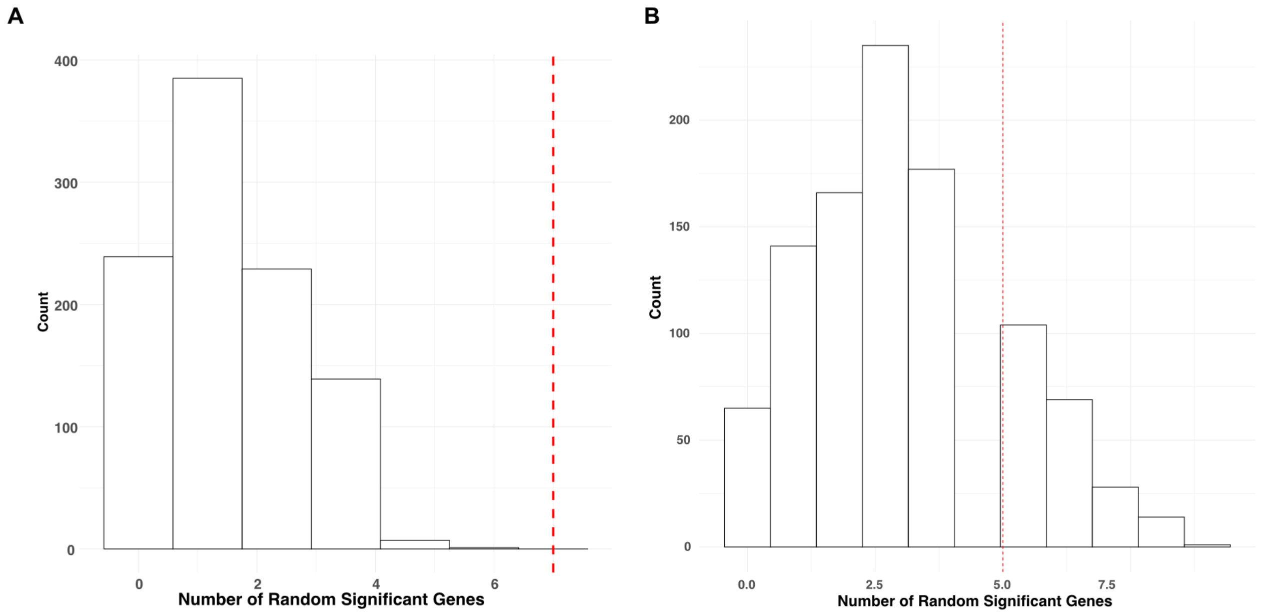


**Supplementary Figure 11:** Histogram showing distribution of number of randomly picked significant genes in Mariathasan dataset (A) and in Uromol dataset (B). Red line indicates the number of significant genes found based on the GENIUS analysis.

**Supplementary Table 1: Table shows the top 50 genes associated to metastatic disease development for every cancer type**

| | Gene | Cancer | DataSource | Attribution |
|---|---|---|---|---|
| 1 | TP53 | UCEC | Mutation | 0.382868181637428 |
| 2 | PTEN | UCEC | Mutation | 0.131854600010379 |
| 3 | PIK3CA | UCEC | Mutation | 0.127915627623053 |
| 4 | NBPF9 | UCEC | Methylation | 0.097576713429995 |
| 5 | FGF11 | UCEC | Methylation | 0.0964600342648074 |
| 6 | POTEI | UCEC | Methylation | 0.0906236010121581 |
| 7 | MEP1A | UCEC | Methylation | 0.083046035261942 |
| 8 | PRKACB | UCEC | Methylation | 0.0800968645065834 |
| 9 | RNASE9 | UCEC | Methylation | 0.0784263863745215 |
| 10 | ARID1A | UCEC | Mutation | 0.0758820505074545 |
| 11 | TOP3A | UCEC | Methylation | 0.0754934395462621 |
| 12 | RPS16 | UCEC | Methylation | 0.0746525526555089 |
| 13 | FAM153C | UCEC | Methylation | 0.0735920052295195 |
| 14 | USP35 | UCEC | Methylation | 0.0729063858466382 |
| 15 | CYB561A3 | UCEC | Methylation | 0.0725787507205281 |
| 16 | FKBP15 | UCEC | Methylation | 0.070101354242521 |
| 17 | FTH1P18 | UCEC | Methylation | 0.0698206971083544 |
| 18 | NUFIP1 | UCEC | Methylation | 0.0696191540877447 |
| 19 | LRRC39 | UCEC | Methylation | 0.0694975213897367 |
| 20 | CD180 | UCEC | Methylation | 0.0694846149705525 |
| 21 | RPL26L1 | UCEC | Methylation | 0.0690783267879889 |
| 22 | NDOR1 | UCEC | Methylation | 0.0683962966599236 |
| 23 | SNORA20 | UCEC | Methylation | 0.0680073488883822 |
| 24 | GID4 | UCEC | Methylation | 0.0679755103324513 |
| 25 | DPY19L2P3 | UCEC | Methylation | 0.0672604819695431 |
| 26 | AGPAT4-IT1 | UCEC | Methylation | 0.0665015614859856 |
| 27 | MYO15A | UCEC | Methylation | 0.0663499856371874 |
| 28 | OR2B2 | UCEC | Methylation | 0.0662823819726957 |

| | | | | |
|---|---|---|---|---|
| 29 | CD200R1 | UCEC | Methylation | 0.0661526469532112 |
| 30 | CTDNEP1 | UCEC | Methylation | 0.0659380720475935 |
| 31 | HEXIM2 | UCEC | Methylation | 0.0653582634232147 |
| 32 | IL17RB | UCEC | Methylation | 0.0644056468335523 |
| 33 | SUPT7L | UCEC | Methylation | 0.0639667973994087 |
| 34 | ITIH4 | UCEC | Methylation | 0.0638667946220767 |
| 35 | MYH3 | UCEC | Methylation | 0.0633554462821101 |
| 36 | UBN1 | UCEC | Methylation | 0.0633475214352981 |
| 37 | PPP1R17 | UCEC | Methylation | 0.0625388973822613 |
| 38 | TDRD6 | UCEC | Methylation | 0.0624210509415097 |
| 39 | MIR3671 | UCEC | Methylation | 0.0621278045928936 |
| 40 | MIR590 | UCEC | Methylation | 0.0616362669054721 |
| 41 | TP53 | COAD | Mutation | 0.425126324384425 |
| 42 | NBPF9 | COAD | Methylation | 0.136495689000121 |
| 43 | FGF11 | COAD | Methylation | 0.118545018685747 |
| 44 | APC | COAD | Mutation | 0.112689114848485 |
| 45 | RBMX | COAD | Expression | 0.102455313775893 |
| 46 | FAM84B | COAD | Methylation | 0.10177980525707 |
| 47 | PIK3CA | COAD | Mutation | 0.0970840928334837 |
| 48 | F12 | COAD | Expression | 0.0969105046659154 |
| 49 | FAM153C | COAD | Methylation | 0.0932118457689732 |
| 50 | FABP1 | COAD | Expression | 0.0931375574167134 |
| 51 | RPL26L1 | COAD | Methylation | 0.0930561568518642 |
| 52 | SNORD111B | COAD | Methylation | 0.0928750705723761 |
| 53 | POTEI | COAD | Methylation | 0.0928036248798739 |
| 54 | MRRF | COAD | Expression | 0.0907593897768143 |
| 55 | CHP2 | COAD | Methylation | 0.0903842796998066 |
| 56 | WIPI2 | COAD | Expression | 0.0903451606886801 |
| 57 | RPS16 | COAD | Methylation | 0.0878636620261555 |
| 58 | PRKACB | COAD | Methylation | 0.0876231789725569 |
| 59 | EIF3B | COAD | Expression | 0.0870207857038066 |
| 60 | YWHAZ | COAD | Expression | 0.0861828598729333 |

| | | | | |
|---|---|---|---|---|
| **61** | **JUP** | **COAD** | **Expression** | **0.0860971041054034** |
| **62** | **NMD3** | **COAD** | **Expression** | **0.0850293541653361** |
| **63** | **TMC3** | **COAD** | **Methylation** | **0.0849335984047096** |
| **64** | **SKP1** | **COAD** | **Methylation** | **0.0848350442052058** |
| **65** | **CYB561A3** | **COAD** | **Methylation** | **0.0846108744604781** |
| **66** | **UBE2U** | **COAD** | **Methylation** | **0.0846042643886199** |
| **67** | **POSTN** | **COAD** | **Expression** | **0.0841051926939231** |
| **68** | **SMIM15** | **COAD** | **Expression** | **0.0838989236522977** |
| **69** | **RPS10** | **COAD** | **Expression** | **0.0826300104569653** |
| **70** | **OR4K15** | **COAD** | **Methylation** | **0.0823982566118867** |
| **71** | **WTAP** | **COAD** | **Expression** | **0.0816708769576127** |
| **72** | **PLA2G5** | **COAD** | **Methylation** | **0.0812681326147641** |
| **73** | **GRK6** | **COAD** | **Expression** | **0.0811497980863758** |
| **74** | **COL7A1** | **COAD** | **Expression** | **0.0808556489957939** |
| **75** | **TAOK1** | **COAD** | **Expression** | **0.0805868289710735** |
| **76** | **SLC3A1** | **COAD** | **Expression** | **0.0804545348738376** |
| **77** | **IL17RE** | **COAD** | **Expression** | **0.0803719477164334** |
| **78** | **FTH1P18** | **COAD** | **Methylation** | **0.0802545124996625** |
| **79** | **GID4** | **COAD** | **Methylation** | **0.0800366313826184** |
| **80** | **MIR4786** | **COAD** | **Methylation** | **0.0798250750072179** |
| **81** | **TP53** | **BLCA** | **Mutation** | **0.283024724144084** |
| **82** | **NBPF9** | **BLCA** | **Methylation** | **0.125387229883799** |
| **83** | **RBMX** | **BLCA** | **Expression** | **0.0893599823545985** |
| **84** | **COL7A1** | **BLCA** | **Expression** | **0.0857388462461531** |
| **85** | **RPL26L1** | **BLCA** | **Methylation** | **0.0847478806995396** |
| **86** | **KRT17** | **BLCA** | **Expression** | **0.0834756993381384** |
| **87** | **FAM84B** | **BLCA** | **Methylation** | **0.082056644624091** |
| **88** | **JUP** | **BLCA** | **Expression** | **0.0813937362747419** |
| **89** | **WIPI2** | **BLCA** | **Expression** | **0.0809694811497541** |
| **90** | **PRKACB** | **BLCA** | **Methylation** | **0.0799907056861803** |
| **91** | **TOP3A** | **BLCA** | **Methylation** | **0.0786341657361653** |
| **92** | **EIF3B** | **BLCA** | **Expression** | **0.0781149551365792** |

| | | | | |
|---|---|---|---|---|
| 93 | RPS16 | BLCA | Methylation | 0.078096451077744 |
| 94 | WTAP | BLCA | Expression | 0.0772607183829284 |
| 95 | SKP1 | BLCA | Methylation | 0.0755058465078352 |
| 96 | POTEI | BLCA | Methylation | 0.0752254042586067 |
| 97 | MRRF | BLCA | Expression | 0.0750039767143019 |
| 98 | SNORD111B | BLCA | Methylation | 0.0749298875662924 |
| 99 | FAM153C | BLCA | Methylation | 0.0744861315713206 |
| 100 | F12 | BLCA | Expression | 0.0740914295922199 |
| 101 | CYB561A3 | BLCA | Methylation | 0.0730468019538089 |
| 102 | ITIH4 | BLCA | Methylation | 0.0720740136397203 |
| 103 | GPAA1 | BLCA | Expression | 0.0715250467977883 |
| 104 | NMD3 | BLCA | Expression | 0.0714585184834919 |
| 105 | FTH1P18 | BLCA | Methylation | 0.071455632095265 |
| 106 | ANKS3 | BLCA | Methylation | 0.0713783213897225 |
| 107 | GRK6 | BLCA | Expression | 0.071325779200591 |
| 108 | CHP2 | BLCA | Methylation | 0.0711581933053296 |
| 109 | TRAPPC5 | BLCA | Expression | 0.0709423842118108 |
| 110 | SMIM15 | BLCA | Expression | 0.0705174169844794 |
| 111 | YWHAZ | BLCA | Expression | 0.0704917003436842 |
| 112 | ACP2 | BLCA | Expression | 0.0699935284442238 |
| 113 | TRPT1 | BLCA | Expression | 0.0696829582221007 |
| 114 | RPS10 | BLCA | Expression | 0.0696085392981794 |
| 115 | RAF1 | BLCA | Expression | 0.0688439885573075 |
| 116 | MIR590 | BLCA | Methylation | 0.068602633861773 |
| 117 | LOC105372330 | BLCA | Loss | 0.0685957077668219 |
| 118 | MIR4786 | BLCA | Methylation | 0.0682851336184208 |
| 119 | GNPTG | BLCA | Expression | 0.0679512012698988 |
| 120 | FLVCR1 | BLCA | Expression | 0.0666160054790637 |
| 121 | SLC3A1 | KIRC | Expression | 0.252675493129146 |
| 122 | VHL | KIRC | Mutation | 0.186871020468348 |
| 123 | NBPF9 | KIRC | Methylation | 0.174981916717459 |
| 124 | RBMX | KIRC | Expression | 0.160914920587921 |

| | | | | |
|---|---|---|---|---|
| 125 | YWHAZ | KIRC | Expression | 0.158619304548634 |
| 126 | RPS10 | KIRC | Expression | 0.156298456640312 |
| 127 | PLVAP | KIRC | Expression | 0.153508456159667 |
| 128 | WTAP | KIRC | Expression | 0.153354132879175 |
| 129 | MCAM | KIRC | Expression | 0.151569968797408 |
| 130 | EPAS1 | KIRC | Expression | 0.148692917074507 |
| 131 | G6PC | KIRC | Expression | 0.146390124465653 |
| 132 | JUP | KIRC | Expression | 0.146281388825153 |
| 133 | MRRF | KIRC | Expression | 0.145489425578116 |
| 134 | CTSF | KIRC | Expression | 0.144867297631199 |
| 135 | WIPI2 | KIRC | Expression | 0.140897933867423 |
| 136 | EIF3B | KIRC | Expression | 0.140648416385634 |
| 137 | HDAC5 | KIRC | Expression | 0.140002151787164 |
| 138 | ACP2 | KIRC | Expression | 0.139823429531749 |
| 139 | POSTN | KIRC | Expression | 0.139082725860182 |
| 140 | MEP1A | KIRC | Methylation | 0.139014356689421 |
| 141 | HEG1 | KIRC | Expression | 0.137100119892396 |
| 142 | TRAPPC5 | KIRC | Expression | 0.137033390318226 |
| 143 | SIRPD | KIRC | Methylation | 0.137012982377538 |
| 144 | TRPT1 | KIRC | Expression | 0.13693925740094 |
| 145 | SLC22A11 | KIRC | Expression | 0.135873245317608 |
| 146 | GPAA1 | KIRC | Expression | 0.135688516776321 |
| 147 | TMED4 | KIRC | Expression | 0.133704192683621 |
| 148 | TAOK1 | KIRC | Expression | 0.133656678551597 |
| 149 | TCN2 | KIRC | Expression | 0.133519422726981 |
| 150 | OGG1 | KIRC | Expression | 0.13328369469376 |
| 151 | JAG1 | KIRC | Expression | 0.132994798774922 |
| 152 | SMIM15 | KIRC | Expression | 0.132675975445554 |
| 153 | RPS16 | KIRC | Methylation | 0.132207866943505 |
| 154 | NMD3 | KIRC | Expression | 0.131824865765409 |
| 155 | OR4K15 | KIRC | Methylation | 0.130816314714031 |
| 156 | EIF4ENIF1 | KIRC | Expression | 0.129095078891774 |

| | | | | |
|---|---|---|---|---|
| 157 | ACTR3 | KIRC | Expression | 0.128838012730246 |
| 158 | F12 | KIRC | Expression | 0.128722952133744 |
| 159 | SNORD111B | KIRC | Methylation | 0.128651430930955 |
| 160 | HIF1A | KIRC | Expression | 0.128606308100579 |
| 161 | TP53 | STAD | Mutation | 0.349145426272471 |
| 162 | NBPF9 | STAD | Methylation | 0.141701159578958 |
| 163 | RBMX | STAD | Expression | 0.106367481375171 |
| 164 | FAM84B | STAD | Methylation | 0.0967168229594636 |
| 165 | JUP | STAD | Expression | 0.0963192538635671 |
| 166 | RPL26L1 | STAD | Methylation | 0.0949397691718784 |
| 167 | FGF11 | STAD | Methylation | 0.0939074690384244 |
| 168 | WIPI2 | STAD | Expression | 0.0937349832266084 |
| 169 | F12 | STAD | Expression | 0.0935574327608484 |
| 170 | WTAP | STAD | Expression | 0.0929403079918611 |
| 171 | FAM153C | STAD | Methylation | 0.0927100754122861 |
| 172 | POTEI | STAD | Methylation | 0.0926633071618398 |
| 173 | YWHAZ | STAD | Expression | 0.0916919309306655 |
| 174 | MRRF | STAD | Expression | 0.0916290156177764 |
| 175 | SNORD111B | STAD | Methylation | 0.0908818995165081 |
| 176 | RPS16 | STAD | Methylation | 0.0906909875188031 |
| 177 | PRKACB | STAD | Methylation | 0.0903151577745731 |
| 178 | EIF3B | STAD | Expression | 0.0894964297129637 |
| 179 | POSTN | STAD | Expression | 0.089486499608681 |
| 180 | BPIFB1 | STAD | Expression | 0.0893842085625569 |
| 181 | GRK6 | STAD | Expression | 0.0890152438369816 |
| 182 | OR4K15 | STAD | Methylation | 0.0890119073985743 |
| 183 | FLVCR1 | STAD | Expression | 0.0874979945914406 |
| 184 | TAOK1 | STAD | Expression | 0.0862768972376305 |
| 185 | SMIM15 | STAD | Expression | 0.0862338499095383 |
| 186 | TOP3A | STAD | Methylation | 0.0859928940356323 |
| 187 | NMD3 | STAD | Expression | 0.0853028482657252 |
| 188 | RPS10 | STAD | Expression | 0.0849207298170021 |

| | | | | |
|---|---|---|---|---|
| 189 | IL17RE | STAD | Expression | 0.0843872751786373 |
| 190 | GPAA1 | STAD | Expression | 0.0833499977569325 |
| 191 | SKP1 | STAD | Methylation | 0.0831031073162102 |
| 192 | COL7A1 | STAD | Expression | 0.08269400402757 |
| 193 | ACP2 | STAD | Expression | 0.0825346151385519 |
| 194 | HIF1A | STAD | Expression | 0.0823336901262277 |
| 195 | PRSS2 | STAD | Expression | 0.0816925057661726 |
| 196 | MIR4786 | STAD | Methylation | 0.0814786877394946 |
| 197 | FTH1P18 | STAD | Methylation | 0.0814474230228642 |
| 198 | NDOR1 | STAD | Methylation | 0.0812241084609286 |
| 199 | CHP2 | STAD | Methylation | 0.0809985564307451 |
| 200 | MIR503 | STAD | Methylation | 0.0809465504479027 |
| 201 | TP53 | OV | Mutation | 0.211414088864187 |
| 202 | KLK10 | OV | Expression | 0.0383816071835576 |
| 203 | RPS10 | OV | Expression | 0.035455225192766 |
| 204 | KLK14 | OV | Expression | 0.031635104988199 |
| 205 | NSL1 | OV | Expression | 0.0307064422597902 |
| 206 | TRPT1 | OV | Expression | 0.0292804891481971 |
| 207 | CTU1 | OV | Expression | 0.0276855255889658 |
| 208 | WTAP | OV | Expression | 0.0271757287516754 |
| 209 | RPL18A | OV | Expression | 0.0267239177946433 |
| 210 | JUP | OV | Expression | 0.0264267864684741 |
| 211 | SMIM15 | OV | Expression | 0.0251085555829532 |
| 212 | KLK6 | OV | Expression | 0.0248943131880786 |
| 213 | KRT7 | OV | Expression | 0.0246205527655771 |
| 214 | TMED4 | OV | Expression | 0.024605340149513 |
| 215 | ACTG1 | OV | Expression | 0.0245813923334269 |
| 216 | LRRC46 | OV | Expression | 0.0244269036801201 |
| 217 | PRPF4 | OV | Expression | 0.023964298744293 |
| 218 | NIF3L1 | OV | Expression | 0.0235750915324921 |
| 219 | OGG1 | OV | Expression | 0.0235476089653803 |
| 220 | SCYL1 | OV | Expression | 0.0235199873138303 |

| | | | | |
|---|---|---|---|---|
| 221 | RBMX | OV | Expression | 0.0233176215991743 |
| 222 | RPL28 | OV | Expression | 0.0231376466249558 |
| 223 | ATG4B | OV | Expression | 0.0230975990938833 |
| 224 | MRRF | OV | Expression | 0.0230129957917421 |
| 225 | EPAS1 | OV | Expression | 0.0228279168302114 |
| 226 | FYTTD1 | OV | Expression | 0.0225115993164818 |
| 227 | GPAA1 | OV | Expression | 0.0223225438653421 |
| 228 | AMY1C | OV | Expression | 0.0223100969720284 |
| 229 | DPP3 | OV | Expression | 0.0221957506693199 |
| 230 | EPHB6 | OV | Expression | 0.0221670221790885 |
| 231 | THOC3 | OV | Expression | 0.0221313620639936 |
| 232 | CASP10 | OV | Expression | 0.0220742763253778 |
| 233 | TUBB4A | OV | Expression | 0.0219161134844587 |
| 234 | GRK6 | OV | Expression | 0.0218678737297592 |
| 235 | MCAM | OV | Expression | 0.0218195475269184 |
| 236 | RPL29 | OV | Expression | 0.021765242417678 |
| 237 | KDM3A | OV | Expression | 0.021713043953537 |
| 238 | C3 | OV | Expression | 0.0215304907857296 |
| 239 | ZBTB22 | OV | Expression | 0.0214424868312736 |
| 240 | KRT17 | OV | Expression | 0.0214289498234091 |

**Supplementary Table 2: Summarized results for all output variables, containing top ten events with respect to the predicted variable.**

| Predicted Var. | Gene Name | Data Source | CancerType |
|---|---|---|---|
| Age | ARID1A | Mutation | UCEC |
| Age | SLC13A2 | Methylation | UCEC |
| Age | FBXW7 | Mutation | UCEC |
| Age | RNF5P1 | Methylation | UCEC |
| Age | SPIDR | Methylation | UCEC |
| Age | KNTC1 | Methylation | UCEC |
| Age | TP53 | Mutation | UCEC |
| Age | CSTF3 | Methylation | UCEC |
| Age | CSMD3 | Mutation | UCEC |
| Age | YME1L1 | Methylation | UCEC |
| Age | APC | Mutation | COAD |
| Age | KRAS | Mutation | COAD |
| Age | TP53 | Mutation | COAD |
| Age | GLA | Expression | COAD |
| Age | FBXW7 | Mutation | COAD |
| Age | SPIDR | Methylation | COAD |
| Age | DENND5A | Mutation | COAD |
| Age | SLC13A2 | Methylation | COAD |
| Age | LGALS4 | Expression | COAD |
| Age | OR8D1 | Methylation | COAD |
| Age | GLA | Expression | BLCA |
| Age | SPIDR | Methylation | BLCA |
| Age | PRKAB1 | Methylation | BLCA |

| | | | |
|---|---|---|---|
| Age | TMEM9B | Expression | BLCA |
| Age | TAF13 | Expression | BLCA |
| Age | RBMY1D | Loss | BLCA |
| Age | C19orf44 | Methylation | BLCA |
| Age | CSTF3 | Methylation | BLCA |
| Age | TP53 | Mutation | BLCA |
| Age | POTEJ | Methylation | BLCA |
| Age | GLA | Expression | KIRC |
| Age | ITM2B | Expression | KIRC |
| Age | GSAP | Expression | KIRC |
| Age | TMEM9B | Expression | KIRC |
| Age | TAF13 | Expression | KIRC |
| Age | DDRGK1 | Expression | KIRC |
| Age | RPS8 | Expression | KIRC |
| Age | RPL35A | Expression | KIRC |
| Age | PHC1 | Expression | KIRC |
| Age | TMEM185B | Expression | KIRC |
| Age | GLA | Expression | STAD |
| Age | CSMD3 | Mutation | STAD |
| Age | SPIDR | Methylation | STAD |
| Age | TP53 | Mutation | STAD |
| Age | TAF13 | Expression | STAD |
| Age | LGALS4 | Expression | STAD |
| Age | ITM2B | Expression | STAD |
| Age | TMEM9B | Expression | STAD |
| Age | CSTF3 | Methylation | STAD |
| Age | POTEJ | Methylation | STAD |

| | | | |
|---|---|---|---|
| Age | TP53 | Mutation | OV |
| Age | GLA | Expression | OV |
| Age | RPS8 | Expression | OV |
| Age | DDRGK1 | Expression | OV |
| Age | ACP2 | Expression | OV |
| Age | TMEM185B | Expression | OV |
| Age | SLC7A8 | Expression | OV |
| Age | GSAP | Expression | OV |
| Age | ITM2B | Expression | OV |
| Age | TAF13 | Expression | OV |
| Predicted Var. | Gene Name | Data Source | CancerType |
| CancerType | TP53 | Mutation | UCEC |
| CancerType | SND1-IT1 | Methylation | UCEC |
| CancerType | ZNF641 | Methylation | UCEC |
| CancerType | CTBP1-AS2 | Methylation | UCEC |
| CancerType | MIR4689 | Methylation | UCEC |
| CancerType | MIR557 | Methylation | UCEC |
| CancerType | CLUHP3 | Methylation | UCEC |
| CancerType | NAV2-AS2 | Methylation | UCEC |
| CancerType | TROAP | Methylation | UCEC |
| CancerType | LINC00930 | Methylation | UCEC |
| CancerType | TP53 | Mutation | COAD |
| CancerType | SND1-IT1 | Methylation | COAD |
| CancerType | CTBP1-AS2 | Methylation | COAD |
| CancerType | MIR4689 | Methylation | COAD |
| CancerType | GPR143 | Methylation | COAD |
| CancerType | CLUHP3 | Methylation | COAD |

| | | | |
|---|---|---|---|
| CancerType | LINC00676 | Methylation | COAD |
| CancerType | ZNF641 | Methylation | COAD |
| CancerType | TBL1X | Expression | COAD |
| CancerType | CHRNA9 | Methylation | COAD |
| CancerType | TP53 | Mutation | BLCA |
| CancerType | SND1-IT1 | Methylation | BLCA |
| CancerType | ZNF641 | Methylation | BLCA |
| CancerType | MIR4689 | Methylation | BLCA |
| CancerType | CTBP1-AS2 | Methylation | BLCA |
| CancerType | CLUHP3 | Methylation | BLCA |
| CancerType | MIR138-2 | Methylation | BLCA |
| CancerType | CUZD1 | Methylation | BLCA |
| CancerType | MIR557 | Methylation | BLCA |
| CancerType | TBL1X | Expression | BLCA |
| CancerType | SND1-IT1 | Methylation | KIRC |
| CancerType | MIR4689 | Methylation | KIRC |
| CancerType | TBL1X | Expression | KIRC |
| CancerType | CTBP1-AS2 | Methylation | KIRC |
| CancerType | CHRNA9 | Methylation | KIRC |
| CancerType | CLUHP3 | Methylation | KIRC |
| CancerType | MMP3 | Methylation | KIRC |
| CancerType | LIPF | Methylation | KIRC |
| CancerType | TPD52L2 | Expression | KIRC |
| CancerType | OR2D2 | Methylation | KIRC |
| CancerType | TP53 | Mutation | STAD |
| CancerType | SND1-IT1 | Methylation | STAD |
| CancerType | ZNF641 | Methylation | STAD |

| | | | |
|---|---|---|---|
| CancerType | CTBP1-AS2 | Methylation | STAD |
| CancerType | CLUHP3 | Methylation | STAD |
| CancerType | MIR4689 | Methylation | STAD |
| CancerType | GPR143 | Methylation | STAD |
| CancerType | TBL1X | Expression | STAD |
| CancerType | HMCN1 | Mutation | STAD |
| CancerType | FAM197Y2 | Methylation | STAD |
| CancerType | TP53 | Mutation | OV |
| CancerType | CREG1 | Expression | OV |
| CancerType | JSRP1 | Expression | OV |
| CancerType | RPL13 | Expression | OV |
| CancerType | MSI1 | Expression | OV |
| CancerType | LRRN2 | Expression | OV |
| CancerType | LYPD1 | Expression | OV |
| CancerType | DNAJB13 | Expression | OV |
| CancerType | SIKE1 | Expression | OV |
| CancerType | NUDT16L1 | Expression | OV |
| Predicted Var. | Gene Name | Data Source | CancerType |
| Metastatic | TP53 | Mutation | UCEC |
| Metastatic | PTEN | Mutation | UCEC |
| Metastatic | PIK3CA | Mutation | UCEC |
| Metastatic | NBPF9 | Methylation | UCEC |
| Metastatic | FGF11 | Methylation | UCEC |
| Metastatic | POTEI | Methylation | UCEC |
| Metastatic | MEP1A | Methylation | UCEC |
| Metastatic | PRKACB | Methylation | UCEC |
| Metastatic | RNASE9 | Methylation | UCEC |

| | | | |
|---|---|---|---|
| Metastatic | ARID1A | Mutation | UCEC |
| Metastatic | TP53 | Mutation | COAD |
| Metastatic | NBPF9 | Methylation | COAD |
| Metastatic | FGF11 | Methylation | COAD |
| Metastatic | APC | Mutation | COAD |
| Metastatic | RBMX | Expression | COAD |
| Metastatic | FAM84B | Methylation | COAD |
| Metastatic | PIK3CA | Mutation | COAD |
| Metastatic | F12 | Expression | COAD |
| Metastatic | FAM153C | Methylation | COAD |
| Metastatic | FABP1 | Expression | COAD |
| Metastatic | TP53 | Mutation | BLCA |
| Metastatic | NBPF9 | Methylation | BLCA |
| Metastatic | RBMX | Expression | BLCA |
| Metastatic | COL7A1 | Expression | BLCA |
| Metastatic | RPL26L1 | Methylation | BLCA |
| Metastatic | KRT17 | Expression | BLCA |
| Metastatic | FAM84B | Methylation | BLCA |
| Metastatic | JUP | Expression | BLCA |
| Metastatic | WIPI2 | Expression | BLCA |
| Metastatic | PRKACB | Methylation | BLCA |
| Metastatic | SLC3A1 | Expression | KIRC |
| Metastatic | VHL | Mutation | KIRC |
| Metastatic | NBPF9 | Methylation | KIRC |
| Metastatic | RBMX | Expression | KIRC |
| Metastatic | YWHAZ | Expression | KIRC |
| Metastatic | RPS10 | Expression | KIRC |

| | | | |
|---|---|---|---|
| Metastatic | PLVAP | Expression | KIRC |
| Metastatic | WTAP | Expression | KIRC |
| Metastatic | MCAM | Expression | KIRC |
| Metastatic | EPAS1 | Expression | KIRC |
| Metastatic | TP53 | Mutation | STAD |
| Metastatic | NBPF9 | Methylation | STAD |
| Metastatic | RBMX | Expression | STAD |
| Metastatic | FAM84B | Methylation | STAD |
| Metastatic | JUP | Expression | STAD |
| Metastatic | RPL26L1 | Methylation | STAD |
| Metastatic | FGF11 | Methylation | STAD |
| Metastatic | WIPI2 | Expression | STAD |
| Metastatic | F12 | Expression | STAD |
| Metastatic | WTAP | Expression | STAD |
| Metastatic | TP53 | Mutation | OV |
| Metastatic | KLK10 | Expression | OV |
| Metastatic | RPS10 | Expression | OV |
| Metastatic | KLK14 | Expression | OV |
| Metastatic | NSL1 | Expression | OV |
| Metastatic | TRPT1 | Expression | OV |
| Metastatic | CTU1 | Expression | OV |
| Metastatic | WTAP | Expression | OV |
| Metastatic | RPL18A | Expression | OV |
| Metastatic | JUP | Expression | OV |
| **Predicted Var.** | **Gene Name** | **Data Source** | **CancerType** |
| TP53 | PTEN | Mutation | UCEC |
| TP53 | PIK3CA | Mutation | UCEC |

| | | | |
|---|---|---|---|
| TP53 | ADCY10P1 | Methylation | UCEC |
| TP53 | CXXC4 | Methylation | UCEC |
| TP53 | SNORD53 | Methylation | UCEC |
| TP53 | LRWD1 | Methylation | UCEC |
| TP53 | PIGG | Methylation | UCEC |
| TP53 | Sep/14 | Methylation | UCEC |
| TP53 | TAPT1 | Methylation | UCEC |
| TP53 | PGPEP1L | Methylation | UCEC |
| TP53 | PIK3CA | Mutation | COAD |
| TP53 | ADCY10P1 | Methylation | COAD |
| TP53 | CXXC4 | Methylation | COAD |
| TP53 | SNORD53 | Methylation | COAD |
| TP53 | LRWD1 | Methylation | COAD |
| TP53 | PIGG | Methylation | COAD |
| TP53 | OR2C1 | Methylation | COAD |
| TP53 | ITGB2 | Expression | COAD |
| TP53 | USP15 | Methylation | COAD |
| TP53 | TAPT1 | Methylation | COAD |
| TP53 | ADCY10P1 | Methylation | BLCA |
| TP53 | LRWD1 | Methylation | BLCA |
| TP53 | SNORD53 | Methylation | BLCA |
| TP53 | LINC01425 | Loss | BLCA |
| TP53 | PIGG | Methylation | BLCA |
| TP53 | PPM1F | Loss | BLCA |
| TP53 | CXXC4 | Methylation | BLCA |
| TP53 | LOC107986345 | Loss | BLCA |
| TP53 | USP15 | Methylation | BLCA |

| | | | |
|---|---|---|---|
| TP53 | AURKAPS1 | Methylation | BLCA |
| TP53 | ITGB2 | Expression | KIRC |
| TP53 | FES | Expression | KIRC |
| TP53 | TUFM | Expression | KIRC |
| TP53 | CXXC4 | Methylation | KIRC |
| TP53 | NFU1 | Expression | KIRC |
| TP53 | ADCY10P1 | Methylation | KIRC |
| TP53 | ADI1 | Expression | KIRC |
| TP53 | NUPR1 | Expression | KIRC |
| TP53 | ZNF771 | Expression | KIRC |
| TP53 | ABCC13 | Methylation | KIRC |
| TP53 | ADCY10P1 | Methylation | STAD |
| TP53 | CXXC4 | Methylation | STAD |
| TP53 | LRWD1 | Methylation | STAD |
| TP53 | ITGB2 | Expression | STAD |
| TP53 | SNORD53 | Methylation | STAD |
| TP53 | PIGG | Methylation | STAD |
| TP53 | PIK3CA | Mutation | STAD |
| TP53 | ABCC13 | Methylation | STAD |
| TP53 | OR2C1 | Methylation | STAD |
| TP53 | TAPT1 | Methylation | STAD |
| TP53 | SEC61G | Expression | OV |
| TP53 | NFU1 | Expression | OV |
| TP53 | TUFM | Expression | OV |
| TP53 | RNF181 | Expression | OV |
| TP53 | MEIS1 | Expression | OV |
| TP53 | VOPP1 | Expression | OV |

| | | | |
|---|---|---|---|
| TP53 | BNIP2 | Expression | OV |
| TP53 | ITGB2 | Expression | OV |
| TP53 | NPRL3 | Expression | OV |
| TP53 | RPL18A | Expression | OV |
| Predicted Var. | Gene Name | Data Source | CancerType |
| wGII | TP53 | Mutation | UCEC |
| wGII | PTEN | Mutation | UCEC |
| wGII | MYO15A | Methylation | UCEC |
| wGII | PIK3CA | Mutation | UCEC |
| wGII | MPHOSPH6 | Methylation | UCEC |
| wGII | CXCR5 | Methylation | UCEC |
| wGII | SRRM5 | Methylation | UCEC |
| wGII | SYNGR4 | Methylation | UCEC |
| wGII | ARID1A | Mutation | UCEC |
| wGII | WRAP53 | Methylation | UCEC |
| wGII | TP53 | Mutation | COAD |
| wGII | MYO15A | Methylation | COAD |
| wGII | PIK3CA | Mutation | COAD |
| wGII | MPHOSPH6 | Methylation | COAD |
| wGII | RPL18 | Expression | COAD |
| wGII | CLNS1A | Expression | COAD |
| wGII | SRRM5 | Methylation | COAD |
| wGII | SYNGR4 | Methylation | COAD |
| wGII | CXCR5 | Methylation | COAD |
| wGII | SPDYE3 | Methylation | COAD |
| wGII | TP53 | Mutation | BLCA |
| wGII | MYO15A | Methylation | BLCA |

| | | | |
|---|---|---|---|
| wGII | RPL18 | Expression | BLCA |
| wGII | MPHOSPH6 | Methylation | BLCA |
| wGII | CLNS1A | Expression | BLCA |
| wGII | ZNF787 | Methylation | BLCA |
| wGII | SRRM5 | Methylation | BLCA |
| wGII | SPDYE3 | Methylation | BLCA |
| wGII | WRAP53 | Methylation | BLCA |
| wGII | OR4N4 | Loss | BLCA |
| wGII | RPL18 | Expression | KIRC |
| wGII | GLYATL1 | Expression | KIRC |
| wGII | CLNS1A | Expression | KIRC |
| wGII | NAT8B | Expression | KIRC |
| wGII | TP53 | Mutation | KIRC |
| wGII | B2M | Expression | KIRC |
| wGII | MPHOSPH6 | Methylation | KIRC |
| wGII | TP53 | Expression | KIRC |
| wGII | MYO15A | Methylation | KIRC |
| wGII | EDF1 | Expression | KIRC |
| wGII | TP53 | Mutation | STAD |
| wGII | MYO15A | Methylation | STAD |
| wGII | BCL11A | Loss | STAD |
| wGII | RPL18 | Expression | STAD |
| wGII | CLNS1A | Expression | STAD |
| wGII | MPHOSPH6 | Methylation | STAD |
| wGII | SRRM5 | Methylation | STAD |
| wGII | SYNGR4 | Methylation | STAD |
| wGII | SPDYE3 | Methylation | STAD |

| | | | |
|---|---|---|---|
| wGII | MYO15A | Expression | STAD |
| wGII | TP53 | Mutation | OV |
| wGII | RPL18 | Expression | OV |
| wGII | MYO15A | Expression | OV |
| wGII | CLNS1A | Expression | OV |
| wGII | B2M | Expression | OV |
| wGII | SMG9 | Expression | OV |
| wGII | TP53 | Expression | OV |
| wGII | MYL6 | Expression | OV |
| wGII | KIAA2013 | Expression | OV |
| wGII | HADHA | Expression | OV |

**Supplementary Table 3: Top 10 expressed or methylated genes for every cancer type associated to metastatic disease**

| **BLCA** | **UCEC** | **STAD** | **OV** | **KIRC** | **COAD** |
|---|---|---|---|---|---|
| TOP3A | TOP3A | F12 | SMIM15 | SLC3A1 | F12 |
| RBMX | POTEI | RBMX | WTAP | RBMX | RBMX |
| PRKACB | PRKACB | FAM153C | RPS10 | YWHAZ | POTEI |
| KRT17 | MEP1A | FGF11 | KLK10 | WTAP | SNORD111B |
| WIPI2 | RNASE9 | WIPI2 | KLK14 | PLVAP | FGF11 |
| RPL26L1 | FGF11 | RPL26L1 | RPL18A | RPS10 | RPL26L1 |
| NBPF9 | NBPF9 | NBPF9 | NSL1 | NBPF9 | NBPF9 |
| JUP | RPS16 | JUP | JUP | MCAM | FABP1 |
| FAM84B | FAM153C | FAM84B | CTU1 | EPAS1 | FAM84B |
| COL7A1 | USP35 | WTAP | TRPT1 | G6PC | FAM153C |

**Supplementary Table 4: Full model specifications for every multivariate cox proportional hazard model in late stage immunotherapy treated bladder cancer validation cohort (Mariathasan).**

| | coef | exp(coef) | se(coef) | robust se | z | Pr(>\|z\|) | X2.5.. | X97.5.. | gene |
|---|---|---|---|---|---|---|---|---|---|
| gene | 0.559494093100846 | 1.74978704699739 | 0.268220314622355 | 0.258774184131026 | 2.16209393135428 | 0.030610936712551 | 0.052306012075 2989 | 1.06668217412639 | RBMX |
| StageII | 0.267806177134807 | 1.30709377080162 | 0.219070966816889 | 0.217994408834121 | 1.2285002104737 | 0.219259250426094 | -0.15945501301 1169 | 0.695067367280784 | RBMX |
| StageIII | 0.0676825651826142 | 1.07002559118444 | 0.219280305517437 | 0.21410213529527 | 0.316122793867854 | 0.751909306213455 | -0.35194990900 9238 | 0.487315039374466 | RBMX |
| StageIV | 0.0285329295919724 | 1.02894389297922 | 0.234300457760082 | 0.222906650929909 | 0.128003940093041 | 0.898145854951072 | -0.40835607814 5091 | 0.465421937329036 | RBMX |
| gendermale | -0.0432551488234602 | 0.957667011280682 | 0.193932227780121 | 0.19757897429956 | -0.218925869904956 | 0.826707793984318 | -0.43050282255 2963 | 0.343992524906042 | RBMX |
| log10(Neoantigen.burden.per.MB) | -0.951877035409128 | 0.386015777728205 | 0.190896817329088 | 0.18377172863413 | -5.17967068429886 | 2.22277950143734E-07 | -1.31206300490 869 | -0.591691065909565 | RBMX |
| Baseline.ECOG.Score | 0.627276164508931 | 1.87250323586947 | 0.132798538899173 | 0.134777252269195 | 4.65416940876677 | 3.25289280827679E-06 | 0.363117604126 039 | 0.891434724891823 | RBMX |
| gene1 | 0.248064651915797 | 1.28154278371188 | 0.0848142173457259 | 0.0777375695990335 | 3.19105232123029 | 0.00141755606804821 | 0.095701815256 0153 | 0.400427488575578 | COL7A1 |
| StageII1 | 0.437413500839001 | 1.54869633190686 | 0.22830676493705 | 0.219675923829582 | 1.99117633472812 | 0.0464615046308584 | 0.006856601862 4569 | 0.867970399815545 | COL7A1 |
| StageIII1 | -0.0493530532334315 | 0.951845018407933 | 0.222664592614581 | 0.216423614653819 | -0.228039132016043 | 0.819615818914016 | -0.47353554335 8891 | 0.374829436892028 | COL7A1 |
| StageIV1 | 0.0272045378573319 | 1.02757795986437 | 0.235079668571176 | 0.220792440881526 | 0.123213175907274 | 0.901938292206229 | -0.40554069432 9149 | 0.459949770043812 | COL7A1 |

| | | | | | | | | | |
|---|---|---|---|---|---|---|---|---|---|
| **gendermale1** | **-0.0382369291638801** | **0.962484873138076** | **0.194705786917197** | **0.206966135135067** | **-0.184749689309927** | **0.853425370536614** | **-0.443883100048061** | **0.367409241720301** | **COL7A1** |
| **log10(Neoantigen.burden.per.MB)1** | **-0.905380018935178** | **0.404388180733742** | **0.183271005825525** | **0.179167504606394** | **-5.05326019315931** | **4.34331557393556E-07** | **-1.25654187516363** | **-0.554218162706732** | **COL7A1** |
| **Baseline.ECOG.Score1** | **0.644167455944115** | **1.90440087146485** | **0.132832945210646** | **0.135887345305888** | **4.74045213330252** | **2.13241827517814E-06** | **0.377833153189816** | **0.910501758698414** | **COL7A1** |
| **gene2** | **0.18933958221041** | **1.20845125136097** | **0.06251343761544** | **0.0640524225524072** | **2.95600969745515** | **0.00311647292143238** | **0.0637991408851512** | **0.31488002353567** | **KRT17** |
| **StageII2** | **0.493166668368301** | **1.63749341700292** | **0.233555648244024** | **0.236947362273102** | **2.08133428301212** | **0.0374033192648741** | **0.0287583720812559** | **0.957574964655345** | **KRT17** |
| **StageIII2** | **-0.00500618973989754** | **0.995006320343302** | **0.220503067952036** | **0.209713949544693** | **-0.0238715152271292** | **0.980955095361883** | **-0.416037977903146** | **0.406025598423351** | **KRT17** |
| **StageIV2** | **0.258958292560934** | **1.29557976838695** | **0.24365286969213** | **0.231437852707792** | **1.11891071201688** | **0.263178232585934** | **-0.194651563405624** | **0.712568148527492** | **KRT17** |
| **gendermale2** | **0.0138906480589686** | **1.01398757136661** | **0.195013150315516** | **0.207606527653816** | **0.0669085322891736** | **0.946654520477727** | **-0.39301066909793** | **0.420791965215867** | **KRT17** |
| **log10(Neoantigen.burden.per.MB)2** | **-0.909767583824799** | **0.402617788046686** | **0.182990856718119** | **0.177754524633916** | **-5.11811210262275** | **3.08609163516078E-07** | **-1.25816005019631** | **-0.561375117453286** | **KRT17** |
| **Baseline.ECOG.Score2** | **0.617860708672577** | **1.85495550393112** | **0.13147196586471** | **0.135740942358038** | **4.55176380787803** | **5.31980385035014E-06** | **0.351813350423294** | **0.883908066921859** | **KRT17** |
| **gene3** | **0.453349205613376** | **1.57357359131212** | **0.127666460846534** | **0.141992509205538** | **3.19276846468809** | **0.0014091588650565** | **0.175049001496049** | **0.731649409730703** | **JUP** |
| **StageII3** | **0.481878447245898** | **1.6191129657** | **0.230221080177059** | **0.229189708508021** | **2.10253091372571** | **0.0355067945918114** | **0.032674872942944** | **0.931082021548852** | **JUP** |
| **StageIII3** | **0.28000378270206** | **1.32313481735277** | **0.225058672914148** | **0.225055713286888** | **1.24415318594968** | **0.213443189501456** | **-0.161097309855213** | **0.721104875259333** | **JUP** |
| **StageIV3** | **0.382986091796904** | **1.46665763135433** | **0.249977775702774** | **0.25379939442216** | **1.50901105445453** | **0.13129595782628** | **-0.114451580568605** | **0.880423764162413** | **JUP** |

| | | | | | | | | | |
|---|---|---|---|---|---|---|---|---|---|
| gendermale3 | -0.0506888588431539 | 0.950574387339752 | 0.193325672050777 | 0.199670229143429 | -0.253862877107946 | 0.799601495208294 | -0.442035316749135 | 0.340657599062828 | JUP |
| log10(Neoantigen.burden.per.MB)3 | -1.11716596897271 | 0.327205793219175 | 0.199793088081692 | 0.199077248778809 | -5.61172095669239 | 2.00324286599827E-08 | -1.5073502067205 | -0.726981731224922 | JUP |
| Baseline.ECOG.Score3 | 0.691089863155778 | 1.99588959484549 | 0.135661310078694 | 0.142548950509684 | 4.84808804754286 | 1.2465707484011E-06 | 0.411699054122815 | 0.970480672188741 | JUP |
| gene4 | 0.566108574625293 | 1.76139934341374 | 0.24471189571924 | 0.230624927322394 | 2.45467209983733 | 0.0141013148014024 | 0.114092023136234 | 1.01812512611435 | WIPI2 |
| StageII4 | 0.170930902421145 | 1.18640876821599 | 0.221595397410023 | 0.218451580171787 | 0.78246585484402 | 0.433940844592332 | -0.257226327081423 | 0.599088131923712 | WIPI2 |
| StageIII4 | 0.0465491506512837 | 1.04764957046181 | 0.219143128619492 | 0.216944784963467 | 0.21456681090132 | 0.830105072294335 | -0.378654814510898 | 0.471753115813465 | WIPI2 |
| StageIV4 | -0.0382125815468567 | 0.962508307636443 | 0.236208558237506 | 0.222254181003 | -0.171931890659645 | 0.863491074771667 | -0.473822771726183 | 0.397397608632469 | WIPI2 |
| gendermale4 | -0.0671924120999271 | 0.935015275774728 | 0.193468905038003 | 0.200635919855652 | -0.334897221535749 | 0.737702618097088 | -0.46043158902207 | 0.326046764822216 | WIPI2 |
| log10(Neoantigen.burden.per.MB)4 | -0.959054373925203 | 0.383255130718988 | 0.18721895103827 | 0.19004266226946 | -5.04652146245651 | 4.49926371996352E-07 | -1.33153114749945 | -0.586577600350953 | WIPI2 |
| Baseline.ECOG.Score4 | 0.655147774162481 | 1.92542702472042 | 0.134658591115232 | 0.140027686652127 | 4.67870168983136 | 2.88697103037865E-06 | 0.380698551485853 | 0.929596996839109 | WIPI2 |
| gene5 | 0.0173713306803036 | 1.01752308972372 | 0.243719991795776 | 0.239474460973656 | 0.0725393873303868 | 0.942172661457009 | -0.451989988045204 | 0.486732649405811 | TOP3A |
| StageII5 | 0.232535506598103 | 1.26179524762036 | 0.22442467758214 | 0.226769772421644 | 1.02542549703555 | 0.305162462620692 | -0.211925080130663 | 0.67699609332687 | TOP3A |
| StageIII5 | 0.0626934806394719 | 1.06470043791232 | 0.220711281909312 | 0.214732260021103 | 0.291961164257809 | 0.770316317326707 | -0.358174015320781 | 0.483560976599725 | TOP3A |
| StageIV5 | 0.0246164598094569 | 1.0249219463716 | 0.23945846068397 | 0.225128029293843 | 0.109344269066233 | 0.912929435432534 | -0.416626369516954 | 0.465859289135868 | TOP3A |

| | | | | | | | | | |
|---|---|---|---|---|---|---|---|---|---|
| **gendermale5** | **-0.0795521085024548** | **0.923529894954958** | **0.195277960660865** | **0.201400286607998** | **-0.394995011388904** | **0.692846569599187** | **-0.474289416730176** | **0.315185199725267** | **TOP3A** |
| **log10(Neoantigen.burden.per.MB)5** | **-0.834674897588346** | **0.434015558004184** | **0.187342844802257** | **0.183820600384696** | **-4.54070379403373** | **5.60667547353606E-06** | **-1.19495665395888** | **-0.474393141217813** | **TOP3A** |
| **Baseline.ECOG.Score5** | **0.601425496560268** | **1.8247180766624** | **0.134174827540843** | **0.136670232382078** | **4.40055955183359** | **1.07972089391446E-05** | **0.333556763332676** | **0.869294229787859** | **TOP3A** |
| **gene6** | **0.784574701855012** | **2.19147471237981** | **0.219757092911302** | **0.235672276471329** | **3.32909204935889** | **0.000871296028050443** | **0.322665527816641** | **1.24648387589338** | **EIF3B** |
| **StageII6** | **0.330414150740942** | **1.39154431825129** | **0.219812815775867** | **0.222435932642099** | **1.48543513998065** | **0.13742859287552** | **-0.10555226610 5149** | **0.766380567587034** | **EIF3B** |
| **StageIII6** | **0.0406856420435767** | **1.04152464251969** | **0.218622706709745** | **0.213858739273674** | **0.190245403025179** | **0.84911683427421** | **-0.378469784711965** | **0.459841068799119** | **EIF3B** |
| **StageIV6** | **-0.011161134814454** | **0.988900919570367** | **0.234288529455955** | **0.224361893832972** | **-0.0497461249937699** | **0.960324699507469** | **-0.450902366230279** | **0.428580096601371** | **EIF3B** |
| **gendermale6** | **-0.0743297120045805** | **0.928365550107919** | **0.193164253409837** | **0.195000938234657** | **-0.381176176266059** | **0.703072525667229** | **-0.456524527896028** | **0.307865103886867** | **EIF3B** |
| **log10(Neoantigen.burden.per.MB)6** | **-1.09186747814582** | **0.335589202660936** | **0.197801176822671** | **0.194298944735397** | **-5.61952345975304** | **1.91484878229169E-08** | **-1.47268641206134** | **-0.711048544230307** | **EIF3B** |
| **Baseline.ECOG.Score6** | **0.702187508181362** | **2.01816262960516** | **0.138893262282086** | **0.137685743397489** | **5.09992894583282** | **3.39781010316339E-07** | **0.432328409937661** | **0.972046606425063** | **EIF3B** |
| **gene7** | **0.00542360034126435** | **1.00543833468729** | **0.219449698515511** | **0.213475278545303** | **0.0254062221078839** | **0.979730948186738** | **-0.412980257197185** | **0.423827457879714** | **WTAP** |
| **StageII7** | **0.235990433083037** | **1.26616219684034** | **0.218444026855928** | **0.217553515571927** | **1.08474658505353** | **0.278033909010372** | **-0.190406622148014** | **0.662387488314089** | **WTAP** |
| **StageIII7** | **0.0630446318111132** | **1.06507437436881** | **0.223733924652118** | **0.221827893919818** | **0.284205158770075** | **0.776253162281339** | **-0.371730051038102** | **0.497819314660328** | **WTAP** |
| **StageIV7** | **0.0273247577946422** | **1.02770150264829** | **0.236329230764853** | **0.224112794957758** | **0.121924131104574** | **0.902959104342811** | **-0.411928248797174** | **0.466577764386458** | **WTAP** |

| | | | | | | | | | |
|---|---|---|---|---|---|---|---|---|---|
| **gendermale7** | **-0.0809854042609102** | **0.922207151641349** | **0.194856230232055** | **0.201121626357125** | **-0.402668801599224** | **0.687191888060805** | **-0.475176548432996** | **0.313205739911176** | **WTAP** |
| **log10(Neoantigen.burden.per.MB)7** | **-0.830879202071424** | **0.435666079368617** | **0.179508311563154** | **0.17410487679997** | **-4.77229137599646** | **1.82141738604254E-06** | **-1.17211849013215** | **-0.489639914010699** | **WTAP** |
| **Baseline.ECOG.Score7** | **0.60001816295654** | **1.82215189575565** | **0.132718245388444** | **0.135841151117407** | **4.41705740875197** | **1.00053663099584E-05** | **0.333774399147959** | **0.866261926765122** | **WTAP** |
| **gene8** | **-0.0469390598453981** | **0.954145541595518** | **0.128627926528548** | **0.110387391552579** | **-0.425221206744798** | **0.6706754262015** | **-0.263294371635773** | **0.169416251944977** | **POTEI** |
| **StageII8** | **0.235494702281109** | **1.26553467679249** | **0.21800838124648** | **0.216955173901536** | **1.08545326689461** | **0.277720951235034** | **-0.189729624825524** | **0.660719029387743** | **POTEI** |
| **StageIII8** | **0.0571066714293076** | **1.05876874474727** | **0.220673908848055** | **0.214355919534897** | **0.266410517391896** | **0.789923066125797** | **-0.36302321073 2056** | **0.477236553590671** | **POTEI** |
| **StageIV8** | **0.0290733648727053** | **1.02950012084992** | **0.234637281311251** | **0.223177396479108** | **0.13027020357515** | **0.896352653258079** | **-0.408346294389763** | **0.466493024135173** | **POTEI** |
| **gendermale8** | **-0.0830363168185042** | **0.920317723601474** | **0.193125180788785** | **0.198105841549084** | **-0.419151277877706** | **0.675105576505287** | **-0.471316631381708** | **0.305243997744699** | **POTEI** |
| **log10(Neoantigen.burden.per.MB)8** | **-0.820700434864872** | **0.440123268834236** | **0.181353551047849** | **0.175013476458689** | **-4.68935565118377** | **2.74066687276466E-06** | **-1.16372054553305** | **-0.477680324196693** | **POTEI** |
| **Baseline.ECOG.Score8** | **0.598797698985845** | **1.81992938154288** | **0.131815050881745** | **0.134563780831169** | **4.44991732015265** | **8.59033538556534E-06** | **0.335057534933212** | **0.862537863038478** | **POTEI** |
| **gene9** | **0.559886160218504** | **1.75047321546468** | **0.227208458036799** | **0.217206256394315** | **2.57767050320175** | **0.00994687954688893** | **0.134169720468873** | **0.985602599968134** | **MRRF** |
| **StageII9** | **0.308584441406805** | **1.36149647137116** | **0.220414964446699** | **0.216164947810593** | **1.42754153498185** | **0.153423861959136** | **-0.115091071021938** | **0.732259953835548** | **MRRF** |
| **StageIII9** | **0.0442957403333799** | **1.04529144402126** | **0.219623813249437** | **0.220191351940694** | **0.201169300896569** | **0.840566195108255** | **-0.387271379177565** | **0.475862859844325** | **MRRF** |
| **StageIV9** | **0.0346793111257719** | **1.03528765032765** | **0.234185973221723** | **0.224977004924604** | **0.154146025445552** | **0.877494598865169** | **-0.406267515876142** | **0.475626138127685** | **MRRF** |

| | | | | | | | | | |
|---|---|---|---|---|---|---|---|---|---|
| **gendermale9** | **-0.0775311528654485** | **0.925398195142123** | **0.192543244574096** | **0.194567312228274** | **-0.398479847295653** | **0.690276510525278** | **-0.458876077401626** | **0.303813771670729** | **MRRF** |
| **log10(Neoantigen.burden.per.MB)9** | **-0.977449567365209** | **0.376269526208791** | **0.191428961569024** | **0.185749126197151** | **-5.26220277519778** | **1.42339654347305E-07** | **-1.34151116487141** | **-0.613387969859008** | **MRRF** |
| **Baseline.ECOG.Score9** | **0.606750333474381** | **1.83446031773813** | **0.133440369351698** | **0.137717055653449** | **4.40577480106173** | **1.05406428394645E-05** | **0.336829864336723** | **0.876670802612039** | **MRRF** |

**Supplementary Table 5: Full model specifications for every multivariate cox proportional hazard model in early stage bladder cancer validation cohort (UROMOL).**

| | coef | exp(coef) | se(coef) | robust se | z | Pr(>\|z\|) | X2.5.. | X97.5.. | gene |
|---|---|---|---|---|---|---|---|---|---|
| **gene** | -0.61589169366388 | 0.540159024121206 | 0.265766708640629 | 0.229521895116529 | -2.68336793468523 | 0.00728847552334337 | -1.06574634175566 | -0.166037045572103 | RBMX |
| **GenderM** | -0.0164418712600733 | 0.983692558537684 | 0.355651965422774 | 0.359914955941825 | -0.0456826563848901 | 0.963563187576173 | -0.721862222403371 | 0.688978479883225 | RBMX |
| **Age** | 0.0420438499314014 | 1.04294021060478 | 0.0152340990682515 | 0.015775235982009 | 2.66518041183984 | 0.00769470301845149 | 0.0111249555590434 | 0.0729627443037594 | RBMX |
| **Tumor_stageT1** | 16.2965207172186 | 11953333.0912239 | 3037.53215748829 | 0.623581385862356 | 26.1337510815561 | 1.50804379467 18E-150 | 15.0743236594988 | 17.5187177749384 | RBMX |
| **Tumor_stageTa** | 14.5508411319482 | 2086171.51255808 | 3037.53216303952 | 0.660491177095658 | 22.0303338432646 | 1.47482411313827E-107 | 13.2563022127342 | 15.8453800511621 | RBMX |
| **gene1** | 0.0617657658608262 | 1.06371315760931 | 0.0921740176913862 | 0.108060150361365 | 0.571586895393677 | 0.567601875553698 | -0.150028237011431 | 0.273559768733084 | COL7A1 |
| **GenderM1** | 0.0569922187451008 | 1.05864757275684 | 0.354608689709509 | 0.370675270173584 | 0.153752417091142 | 0.877804953258954 | -0.669517960754778 | 0.78350239824498 | COL7A1 |
| **Age1** | 0.0385186321216978 | 1.03927009198442 | 0.0146674344870526 | 0.0151156879543192 | 2.5482553118392 | 0.0108263205632346 | 0.00889242812968623 | 0.0681448361137094 | COL7A1 |
| **Tumor_stageT11** | 16.2113443466427 | 10977347.0320696 | 2969.99494675764 | 0.635793366017035 | 25.4978192808138 | 2.08414294668 65E-143 | 14.9652122476398 | 17.4574764456456 | COL7A1 |
| **Tumor_stageTa1** | 14.3680377424672 | 1737638.80754937 | 2969.99495271207 | 0.672730708202835 | 21.3577848718258 | 3.30097892808737E-101 | 13.0495097830955 | 15.6865657018388 | COL7A1 |
| **gene2** | -0.221135604996578 | 0.801607970873444 | 0.0548982140353723 | 0.0591736142676817 | -3.73706436108896 | 0.000186181279135306 | -0.3371137577963 | -0.105157452196857 | KRT17 |
| **GenderM2** | -0.108784041035554 | 0.89692409538307 | 0.360156717278922 | 0.354215856692055 | -0.30711228472792 | 0.758757907405708 | -0.803034362904984 | 0.585466280833875 | KRT17 |
| **Age2** | 0.0441191883421347 | 1.04510691202561 | 0.0153811655529445 | 0.0161259230437092 | 2.73591708347795 | 0.00622066982886471 | 0.0125129599590002 | 0.0757254167252692 | KRT17 |
| **Tumor_stageT12** | 16.3294534428451 | 12353542.7573608 | 3389.15564498755 | 0.627855918970074 | 26.0082814376135 | 3.99142338983 23E-149 | 15.0988784541835 | 17.5600284315068 | KRT17 |
| **Tumor_stageTa2** | 14.5517997595632 | 2088172.33304734 | 3389.1556495348 | 0.661978556745285 | 21.982282675604 | 4.25520158924 38E-107 | 13.2543456298046 | 15.8492538893218 | KRT17 |
| **gene3** | -0.0126294783757803 | 0.987449938802157 | 0.178627282039849 | 0.192004890018746 | -0.0657768579464159 | 0.947555481009674 | -0.388952147668097 | 0.363693190916537 | JUP |
| **GenderM3** | 0.038863667070909 | 1.03962873835703 | 0.354423806233937 | 0.357030251221709 | 0.108852588647385 | 0.913319411883073 | -0.660902766714928 | 0.738630100856746 | JUP |

| | | | | | | | | | |
|---|---|---|---|---|---|---|---|---|---|
| **Age3** | 0.0393677515746401 | 1.04015293120132 | 0.0147341161840062 | 0.0151607443065491 | 2.59668989718625 | 0.00941268599056879 | 0.00965323875498311 | 0.0690822643942971 | JUP |
| **Tumor_stageT13** | 16.2297191889651 | 11180918.6231398 | 2949.91908851554 | 0.645719453393501 | 25.134319716824 | 2.09742793126812E-139 | 14.9641323161969 | 17.4953060617332 | JUP |
| **Tumor_stageTa3** | 14.3813442434741 | 1760915.22044929 | 2949.91909407296 | 0.67295324127009 | 21.3704955433926 | 2.51450037353388E-101 | 13.0623801273052 | 15.7003083596429 | JUP |
| **gene4** | 0.293680123998319 | 1.34135476783553 | 0.290244530321498 | 0.314888364662432 | 0.932648382588385 | 0.351001543906311 | -0.323489729890763 | 0.9108499778874 | WIPI2 |
| **GenderM4** | 0.0329323708425953 | 1.03348064345272 | 0.353874839658766 | 0.357899707939811 | 0.0920156404490098 | 0.926685613029742 | -0.668538166796838 | 0.734402908482029 | WIPI2 |
| **Age4** | 0.041697481522205 | 1.04257903161725 | 0.0150297929341513 | 0.016328460432488 | 2.55366889576689 | 0.0106594567214746 | 0.00969428715154125 | 0.0737006758928688 | WIPI2 |
| **Tumor_stageT14** | 16.3363274350577 | 12438753.4473528 | 2993.19015241119 | 0.633989650672784 | 25.7674985983158 | 2.05243273024 32E-146 | 15.0937305531679 | 17.5789243169475 | WIPI2 |
| **Tumor_stageTa4** | 14.5132878239893 | 2009281.64174609 | 2993.19015904713 | 0.677639290451491 | 21.4174237363356 | 9.1934870874493E-102 | 13.1851392201951 | 15.8414364277835 | WIPI2 |
| **gene5** | 0.656931023850328 | 1.92886360509098 | 0.230516222010314 | 0.243355711921143 | 2.69946827491438 | 0.00694503770936325 | 0.179962593052784 | 1.13389945464787 | TOP3A |
| **GenderM5** | 0.0511037458792088 | 1.05243207311036 | 0.354359443268429 | 0.361698595003583 | 0.14128820676979 | 0.887642265791299 | -0.657812473586553 | 0.760019965344971 | TOP3A |
| **Age5** | 0.0426440117562642 | 1.04356633137289 | 0.0153540406944314 | 0.015995916745884 | 2.66593108939737 | 0.0076775428438379 | 0.0112925910346304 | 0.0739954324778979 | TOP3A |
| **Tumor_stageT15** | 16.5482065227299 | 15374279.71462 | 3076.70329586956 | 0.614393462076772 | 26.9342164983228 | 8.73242524795292E-160 | 15.3440174647225 | 17.7523955807372 | TOP3A |
| **Tumor_stageTa5** | 14.8439819975791 | 2796787.50710188 | 3076.70330324857 | 0.660468572256481 | 22.4749255621124 | 7.30259137699807E-112 | 13.5494873830358 | 16.1384766121224 | TOP3A |
| **gene6** | -0.406909300540237 | 0.665704566607983 | 0.271444825590996 | 0.285280849927737 | -1.42634635533128 | 0.153768390173301 | -0.966049491877577 | 0.152230890797102 | EIF3B |
| **GenderM6** | 0.0208011750803086 | 1.02101902742893 | 0.354375765005089 | 0.363464724996336 | 0.0572302445045205 | 0.954361786004794 | -0.691576595563265 | 0.733178945723882 | EIF3B |
| **Age6** | 0.0412096592868578 | 1.04207056241489 | 0.014851151838903 | 0.0152044018714799 | 2.71037687869576 | 0.00672067948564237 | 0.0114095792122837 | 0.0710097393614318 | EIF3B |
| **Tumor_stageT16** | 16.1470540703879 | 10293817.9130321 | 2950.98519019172 | 0.650978787952175 | 24.8042706908819 | 8.06250578015971E-136 | 14.8711590913021 | 17.4229490494737 | EIF3B |
| **Tumor_stageTa6** | 14.2798133263269 | 1590904.57310201 | 2950.98519634199 | 0.688347655111905 | 20.7450598840283 | 1.35855174919891E-95 | 12.930676713465 | 15.6289499391888 | EIF3B |
| **gene7** | -1.02549040771817 | 0.358620551988238 | 0.253339960538938 | 0.24968491224443 | -4.10713806653349 | 4.0059179574228E-05 | -1.5148638432003 | -0.536116972236046 | WTAP |

| | | | | | | | | | |
|---|---|---|---|---|---|---|---|---|---|
| **GenderM7** | 0.0251313329506931 | 1.02544978702748 | 0.354980391661299 | 0.376672398735803 | 0.0667193376393905 | 0.946805139402961 | -0.713133002541791 | 0.763395668443177 | WTAP |
| **Age7** | 0.0457414203129376 | 1.04680369378591 | 0.0152914151132206 | 0.0157153736466855 | 2.91061614832077 | 0.00360716868497588 | 0.014939853961844 | 0.0765429866640311 | WTAP |
| **Tumor_stageT17** | 16.3524404295754 | 12640802.4492311 | 3490.55960741173 | 0.644693733744171 | 25.3646647604371 | 6.19211843269522E-142 | 15.0888639303782 | 17.6160169287727 | WTAP |
| **Tumor_stageTa7** | 14.6635599440273 | 2335087.55233125 | 3490.55961090892 | 0.663463057099349 | 22.1015470072081 | 3.05421996034886E-108 | 13.3631962470397 | 15.9639236410148 | WTAP |
| **gene8** | 0.0929256476091544 | 1.09738013939766 | 0.0475272783926474 | 0.046552922773896 | 1.99612918098585 | 0.0459198634684553 | 0.00168359559724381 | 0.184167699621065 | POTEI |
| **GenderM8** | -0.0236041713430397 | 0.976672228111686 | 0.355707319019434 | 0.358375930463084 | -0.0658642764109158 | 0.947485882093281 | -0.726008087976714 | 0.678799745290635 | POTEI |
| **Age8** | 0.0364122448469951 | 1.03708329062151 | 0.0149057149888517 | 0.0155282643926597 | 2.34490113809547 | 0.0190321252972812 | 0.00597740589496634 | 0.0668470837990239 | POTEI |
| **Tumor_stageT18** | 16.0728539843721 | 9557664.7709943 | 3044.54326934654 | 0.631259533602414 | 25.4615623666688 | 5.25716180150985E-143 | 14.8356080336138 | 17.3100999351304 | POTEI |
| **Tumor_stageTa8** | 14.3651108037523 | 1732560.28114455 | 3044.5432730494 | 0.660818566808989 | 21.7383583411096 | 8.90264383475852E-105 | 13.0699302124913 | 15.6602913950133 | POTEI |
| **gene9** | -0.140330582189782 | 0.869070888548136 | 0.193323080726665 | 0.171499077490082 | -0.818258525022662 | 0.41320958321931 | -0.476462597452186 | 0.195801433072623 | MRRF |
| **GenderM9** | 0.0680696578934979 | 1.07043987046832 | 0.357043605237894 | 0.363887210741656 | 0.18706251795649 | 0.851611615702937 | -0.645136169594884 | 0.781275485381879 | MRRF |
| **Age9** | 0.0384763675179579 | 1.03922616857401 | 0.0148126911662582 | 0.0153884753220799 | 2.50033656438664 | 0.0124075368107955 | 0.00831551010969781 | 0.0686372249262179 | MRRF |
| **Tumor_stageT19** | 16.1624765990318 | 10453805.1473768 | 2976.02477880558 | 0.628348105760576 | 25.7221696872439 | 6.60460089261918E-146 | 14.9309369419871 | 17.3940162560765 | MRRF |
| **Tumor_stageTa9** | 14.3088894850265 | 1637841.02610718 | 2976.02478519593 | 0.673100449354207 | 21.2581784765631 | 2.7695904740614E-100 | 12.9896368463145 | 15.6281421237385 | MRRF |
